# Adaptive time scales in recurrent neural networks

**DOI:** 10.1101/800540

**Authors:** Silvan C. Quax, Michele D’Asaro, Marcel A. J. van Gerven

## Abstract

Recurrent neural network models have become widely used in computational neuroscience to model the dynamics of neural populations as well as in machine learning applications to model data with temporal dependencies. The different variants of RNNs commonly used in these scientific fields can be derived as discrete time approximations of the instantaneous firing rate of a population of neurons. The time constants of the neuronal process are generally ignored in these approximations, while learning these time constants could possibly inform us about the time scales underlying temporal processes and enhance the expressive capacity of the network. To investigate the potential of adaptive time constants, we compare the standard Elman approximation to a more lenient one that still accounts for the time scales at which processes unfold. We show that such a model with adaptive time scales performs better on predicting temporal data, increasing the memory capacity of recurrent neural networks, and allows recovery of the time scales at which the underlying processes unfold.

## Introduction

Recurrent neural network (RNN) models have become widely used in computational neuroscience to model the dynamics of neural populations as well as in machine learning applications to model data with temporal dependencies. The different variants of RNNs commonly used in these scientific fields can be derived as discrete time approximations of the instantaneous firing rate of a population of neurons [1]. Since such models can mimic the dynamic properties of real neural populations, they are ideally suited to explain neuronal population data such as extracellular multi-unit activity (MUA), functional magnetic resonance imaging (fMRI) or magnetoencephalography (MEG) data. Similarly, it has turned out that these biologically inspired network enable machines to perform tasks depending on sequential data [2, 3]. The interplay between both fields has led to recent advances, enabling such RNNs to solve a wide variety of cognitive tasks [4, 5, 6].

While RNNs have been successfully applied in both computational neuroscience and machine learning, further improvements could be possible through more biologically inspired RNNs. A common assumption that the popular discrete RNN definitions make, when derived from their continuous counterparts, is that of ignoring certain time scales at which the population activity unfolds. The time scale at which processes unfold is an important aspect of modelling dynamic neuronal activity. Some neural responses, like retinal responses to a flashing head light, act at very short time scales, while others, like maintaining the concept of a car in mind, can take very long. Understanding these time scales can give us valuable insights about the nature of the underlying processes and the kind of information that is processed by a neuronal population.

An interesting example of this is the temporal hierarchy found in the brain [7, 8]. Much like the spatial hierarchies found in the visual cortex that ensure an increase in receptive field size along the visual pathway, there is evidence for a temporal hierarchy with lower visual areas responding at shorter time scales and higher visual areas at longer time scales [9]. A tempting explanation for the emergence of such a hierarchy is the hierarchical causal structure of the outside world that shapes the representations of the brain [10]. For example, a car passing by leads to changes in neuronal activity on a short time scale in the retina, where the amount of light reaching a receptor could suddenly change, while on the other hand, neurons in V4 or IT, encoding the concept ‘car’ would change activity at a much longer time scale. As for the spatial receptive field, one can define a property of the neurons (or of the coding unit more in general, be it an artificial neuron or a population of neurons in a voxel) that defines the region of time of interest for a stimulus to trigger its activity. The idea of such temporal receptive windows of a neuron that represents the length of time before a response during which sensory information may affect that response has been suggested [7], emphasizing the important role of time scales in the neural responses of different cortical areas.

While a hierarchy of time scales can thus be important for processing temporal information, there is no specific time scale parameter in the RNNs commonly used for modelling. Often either one or multiple implicit assumptions about the time scales at which the networks operate are made in the definition of the RNNs used. In computational neuroscience models it is often assumed that either the firing rate closely tracks the intrinsic current, or the intrinsic current closely tracks the firing rates. Models used for artificial intelligence problems on the other hand, tend to ignore time scales both for the intrinsic currents and firing rates (see Methods for further details). By allowing our models to learn the time scales at which processes operate, a closer relation to the temporal hierarchies of our natural world could be achieved, allowing a model to perform better. At the same time this would help in interpreting the role different neurons of the model perform by linking their time scales to the underlying processes.

Here we investigate whether a more biologically plausible model with learnable intrinsic time parameters is able to recover the time scales of the processes underlying a given dataset and whether the flexibility of learning these time scales is beneficial for the performance of the model. We find an improvement in performance compared to the commonly used approximations for RNN models and an increase in the memory capacity of the RNN models. At the same time we can recover the time scales of the underlying processes from the data, opening interesting opportunities to improve our understanding of the temporal hierarchies in the brain.

## Methods

### Synaptic coupling between neurons

Deriving the equations governing the communication between neurons requires specification of a mechanism for synaptic transmission, referring to how incoming spikes generate synaptic currents through the release of neurotransmitters in the synaptic cleft [1]. That is, we need to make explicit how the synaptic current *I* arises. Consider a presynaptic neuron indexed by *k* whose spike times are given by {*t*_*i*,…,*N*_}, with N the number of spikes. Formally, this spike train is fully described by the neural response function

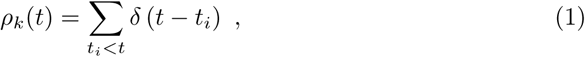

where the idealized action potentials are given by Dirac *δ* functions. The total synaptic current received by a neuron *n* is modeled as

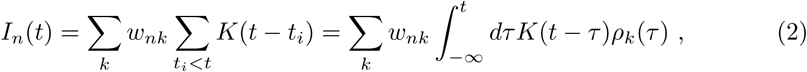

where *w*_*nk*_ is a measure of the efficacy of the synapse from neuron *k* to neuron *n* and *K*(·) determines how presynaptic spikes are transformed into synaptic currents.

A common choice for *K* is simply *K*(*s*) = *qδ*(*s*), where *q* is the charge injected via a synapse with strength *w*_*nk*_ = 1. A more realistic choice for *K* is to assume that the synaptic current has a finite duration, as modeled by the exponential kernel

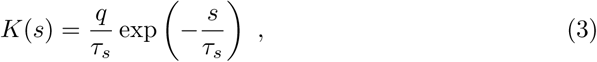

where *τ*_*s*_ is the time constant. A detailed description of why this kernel function is a suitable approximation can be found in [11]. By coupling multiple spiking neurons via the mechanism outlined here we can build networks of spiking neurons.

### From spiking to rate-based models

An alternative to spiking neuron models is to assume that neural coding is driven by the rate at which neurons fire, rather than the exact arrival times of individual spikes. A mean firing rate over a temporal window of length *T* can simply be computed from individual spike times as

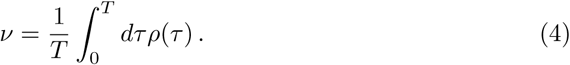

The use of mean firing rates to quantity neural responses dates back to the work of Adrian [12], who showed that the firing rate of muscular stretch receptors is related to the force applied to the muscle. However, a problem associated with the use of mean firing rates is that it severly limits the speed at which information can be processed given the requirement to integrate over time. This is unrealistic given the very rapid response times observed in visual detection tasks [13].

An alternative formulation is provided by assuming that the rate code captures a population average by counting the number of spikes produced by a population consisting of *M* neurons over a short time interval Δ*t*. That is, we interpret the firing rate as a population activity, given by [14]:

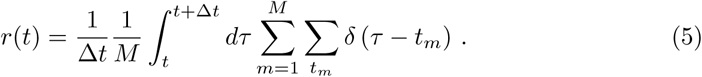

Under this interpretation, we view artificial neurons as models of neuronal populations that are influenced by presynaptic populations in a homogeneous manner and collectively produce a firing rate as output. The population activity may vary rapidly and can reflect changes in the stimulus conditions nearly instantaneously [15].

To compute the postsynaptic current induced by the presynaptic firing rates, we replace the neural response function in Eq. (2) by the firing rate, to obtain

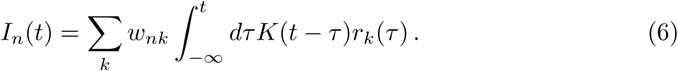

Using an exponential kernel (3) with *q* = 1 and taking the time-derivative of Eq. (6) yields

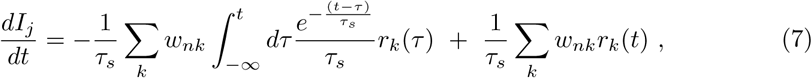

where the Leibniz rule is applied to calculate the derivative of the integral. Substituting the first term on the right hand side for Eq. (6) and using a compact vector notation this simplifies to

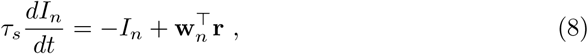

with **w**_*n*_ = (*w*_*n*1_, …, *w*_*nK*_)^*T*^ and **r** = (*r*_1_, …, *r*_*K*_)^*T*^. The kernel time constant, *τ*_*s*_, thus determines the time scale of the differential process.

To complete the model, we assume that the synaptic input *I*_*n*_ directly influences the firing rate *r*_*n*_ of a neuron. Due to the membrane capacitance and resistance the firing rate does not follow the current instantaneously, but with a delay determined by a time constant *τ*_*r*_. That is,

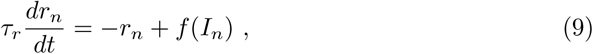

where *f* is an activation function (a static nonlinearity) which translates currents into firing rates. Equations (8) and (9) together define the firing-rate model.

### Discrete approximation of the recurrent dynamics

As mentioned before, firing-rate models provide a biological counterpart for recurrent neural networks, where RNN units reflect the average activity of a population of neurons. The equations for the standard RNN follow from the continuous equations through discretization.

We use the forward Euler method to numerically approximate the solution to the differential equations (8) and (9) and further generalize by assuming that at each point in time, the firing rates are influenced by sensory inputs **x**. We obtain

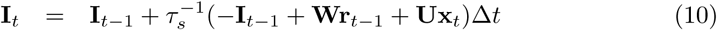

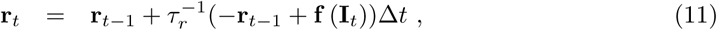

where **W** and **U** are matrices whose values represent synaptic weights, Δ*t* indicates discrete time steps. We can rewrite the last equations, defining *α*_*s*_ = Δ*t/τ*_*s*_ and *α*_*r*_ = Δ*t/τ*_*r*_, as ^1^:

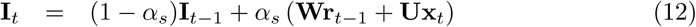

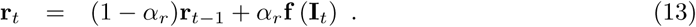

These are going to be the key equations of the model under investigation.

### Time scale assumptions

In literature, several variants of Eqs. (12) and (13) are commonly used to model recurrent dynamics. These variants have implicitly different underlying time scale approximations. Here we review the two approximations that are at the basis of commonly used rate-based RNN models in computational neuroscience and artificial intelligence (AI), derived from our set of equations above. Typically, either the process of integrating the presynaptic current, or the process of generating postsynaptic rate activity, are modeled in literature as instantaneous; a choice which resides in the assumption that one of the two processes is much faster than the other [1]. We will refer to these complexity reduction choices as *extreme cases* for the time constants. Despite the popularity of such choices, it is often neglected in literature why this approximation is made. Furthermore, the implicit time constants used in these models are typically not motivated.

The first approximation assumes that the time constant for the firing rate *τ*_*r*_ is much larger than that of the current *τ*_*s*_. In this case, the current closely tracks the firing rates and we can assume that, for the *n*th-neuron, 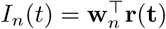. By using this substitution, generalizing to multiple neurons and adding the external input again, the equations describing the dynamics at the network level are then given by

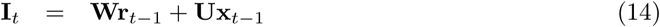

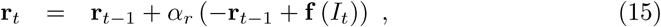

with **W** = [**w**_1_, …, **w**_*K*_]^*T*^ and **f** a vector-valued activation function.

The second approximation on the other hand, assumes that the time constant for the firing rate *τ*_*r*_ is much smaller then that of the current *τ*_*s*_. The firing rate then closely tracks the current and we can assume *r*_*n*_ = *f* (*I*_*n*_). The model is then fully described by Eq. (8). Generalizing to multiple neurons, we obtain

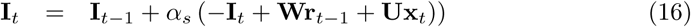

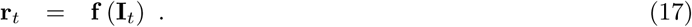

With respect to our discrete-time Eqs. (12) and (13), these two extreme cases correspond respectively to the choices *α*_*s*_ = 1 (Eqs. (14) and (15)) and *α*_*r*_ = 1 (Eqs. (16) and (17)). Studies in computational neuroscience that implement RNNs as modeling tool often use the extreme case of Eqs. (16) and (17) [16, 17, 18].

Studies in AI, on the other hand, tend to ignore both processes, equivalent to assuming both *τ*_*r*_ = Δ*t* and *τ*_*s*_ = Δ*t*. These models, referred to as Elman networks [19], are determined by

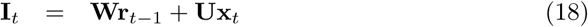

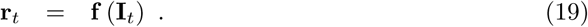

Setting the time constants *α*_*r*_ = 1 and *α*_*s*_ = 1, as in Eqs. (18) and (19), means that the synaptic current and the firing rate both follow instantaneously the presynaptic firing rates. Thus, every neuron acts as a non-linear filter that carries no memory of its previous states internally. The activity of a neuron is influenced by the recent firing rate history only through the recurrent weights.

### Optimizing time constants

While deliberately disregarding one or both of the dynamical processes simplifies the equations describing an RNN, such a simplification could prevent us from achieving optimal performance and gaining valuable insight into the dynamics of the underlying process. We aim here to show that, from a functional perspective such enrichment of the internal dynamics increases the performance of RNNs commonly used in the literature. At the same time it could provide us with valuable insight into the dynamics of the underlying data that the RNN tries to explain.

Previous work has experimented with manually setting the time constants of the dynamic process to enhance the expressiveness of RNNs [18, 20]. While it is possible to come up with time constant values deemed biologically relevant, a much more interesting approach would be to optimize the time constants to adapt to the process at hand. The idea that time constants can be inferred from the data and not set a priori, with the aim to better describe the observed data, has been suggested previously in the continuous time regime [21]. Other work has optimized networks with a single time constant in the discrete approximation regime through numerical integration [22], though limited by the computationally intensive integration process.

Here, instead, we optimize the time constants of the RNN using the backpropagation-through-time (BPTT) algorithm, alongside the other parameters of the network.

### Adaptive time scales recurrent neural network

To investigate the potentially beneficial role time constants can play in the performance and interpretability of RNNs, we developed an RNN model which can adapt the time constants at which dynamic processes unfold. The model was developed using the Chainer package for automatic differentiation [23]. The model consisted of an RNN with hidden units that generate a firing rate according to Eqs. (12) and (13), referred to as adaptive recurrent units (ARUs) throughout the next sections (Fig. 1). The output units transform the hidden units activity in a readout layer. All layers were fully connected. The non-linear transformation of the synaptic currents into firing rate is modelled by a sigmoid function, unless otherwise specified.

**Figure 1:**
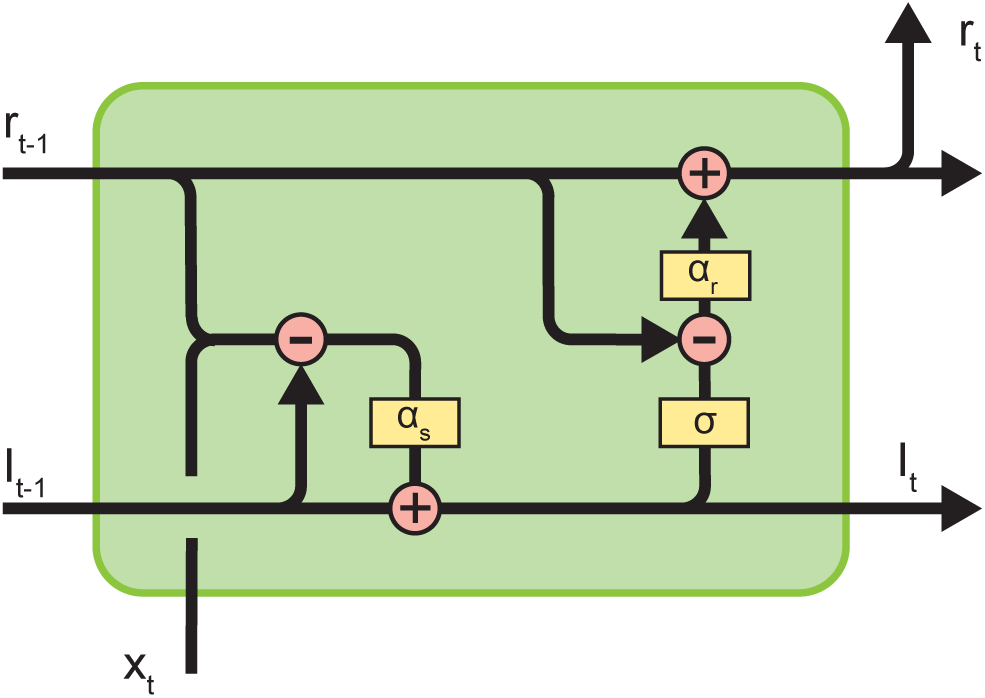
Diagram of the ARU implementation with internal state and time constants as described by Eqs. (12) and (13).

The complete model is then given by

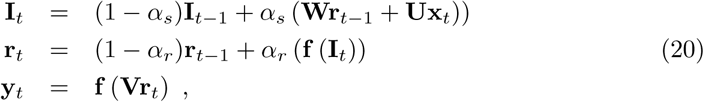

where **x** is the input signal, **W, U, V** weight matrices, **I, r** are the *N* -dimensional state variables referring to the postsynaptic current and the firing rates, with *N* the number of hidden (recurrent) units, and **f** a sigmoid function. The initial states of the synaptic current and firing rate were also learned through the BPTT algorithm [24]. We always used an additional input with a fixed value of one as a bias term.

### Simulations

To investigate whether the explicit modelling of time scales benefits performance and leads to recovery of the relevant time scales of the dynamics of the data, several simulations were performed. Artificial training data was created that has temporal dynamics reflective of a certain time scale by using our previously described model with a certain set of time constants as a generative model to produce data. A random input pattern that consisted of a two-valued signal was generated by drawing samples from a white noise distribution between 0 and 1, and applying a smoothing Savitzky-Golay filter [25].

The data consisted of a series of 500 samples (400 for training and 100 for validation sets) of input-output pairs unfolding over 20 time steps. Target networks are structured as explained in Fig. 1, with two units in the input layers, 10 units in the hidden layer and two units in the output layer (unless stated otherwise). The data generated this way reflects the time scales of the underlying generating process, thus expressing slower dynamics for *α*_*s*_, *α*_*r*_ close to one and expressing fast dynamics for *α*_*s*_, *α*_*r*_ close to zero.

In the simulations, this data was used to optimize networks to learn to produce the output data from the input data. The mean-squared-error between the outputs of the network and the training data set was minimized via BPTT. During training, the data was divided into minibatches. For optimization, the Adam [26] optimizer was used with a learning rate of 0.001.

#### Influence of time constants

We first ran a series of grid search experiments over the *α*_*s*_, *α*_*r*_ parameter space, where the time constants of the trained network were fixed to a certain value, in order to visualize the loss landscape and check the relevance of the time constants for performance. To avoid confusion, the time constants of the trained network will from here on be indicated as 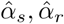. During the grid search the 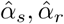 values tested ranged from 0.001 to 1.3 to cover a biologically plausible range.^2^ For each pair of time constants 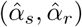 a network was trained over the generated data, minimizing the mean squared error between the outputs of the network and the training data set. Six different combinations of time constants were used to generate data, that is (*α*_*s*_, *α*_*r*_) *∈* {(0.34, 0.68), (0.34, 1.0), (0.68, 0.34), (0.68, 0.68), (0.89, 0.89), (0.14, 0.14)}.

#### Learning optimal time constants

Next, we investigated the idea of learning the best time constants by BPTT and check whether the time constants are recovered correctly and lead to an improvement in performance. For two combinations of time constants (*α*_*s*_ = 0.34, *α*_*r*_ = 0.68 and *α*_*s*_ = 0.68, *α*_*r*_ = 0.34) we trained 20 repetitions of networks with optimizable time constants. The performance was compared with the standard Elman network with fixed time constants of 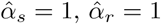.

To investigate the influence of network size on the ability to recover the time constants, networks with different numbers of hidden units were trained (5, 10, 30 and 100 units). A grid search over fixed time constants in the range 0.001 to 1.3 was performed to identify the loss landscape. On top of that, networks with learnable time constants were trained, with 20 repetitions per network size to investigate the distribution of learned time constants.

The influence of the activation function was compared by performing simulations with networks using the sigmoid and rectified linear unit activation functions.

#### Learning individual time constants

In the previous simulations we only used models with time constants that were shared across all units in the network. To investigate whether having individual time constants per unit can help in learning processes with a range of underlying time scales, data was generated with three different distributions of time constants. Time constants were drawn from a Gaussian distribution truncated between 0 and 1. The mean for all three distributions was 0.5, the standard deviation varied from 0.1 to 0.3. The networks were extended with individual time constants per unit and 20 repetitions were trained per data set. The standard deviation of the learned time constants was compared with the standard deviation of the time constants used to generate the data, to infer whether the original distribution was recovered. To investigate whether there was also a benefit in performance the network with individual time constants was compared with an Elman network and a network with global time constants.

#### Testing memory capacity

The memory capacity of the resulting networks was investigated by generating white noise data where the network had to remember the input *N* steps back, with *N* ∈ {5, 10, 20, 30, 40}. To generate a data set with a common underlying data distribution, one data set was generated by low-pass filtering a white noise process to 20 Hz. For every memory length sequences are generated by iterating through windows of 250 ms of data using time-step Δ*t* = 100*/N* ms. The Elman network, the network with global time constants and the network with individual time constants were all trained for 20 repetitions on every memory length task.

## Results

### Optimal time constants improve performance

To investigate whether explicitly modelling time constants in an RNN benefits performance compared to the commonly used approximations (see Methods), we performed several simulations. Artificial data with several combinations of time constants, (*α*_*s*_, *α*_*r*_), was created by using an RNN as generative model (see Methods for further details). A grid search for parameters 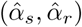 over the range 0.001 to 1.3 was performed to identify the combination of time constants resulting in the best performance. In these simulations, we were interested in seeing whether a network trained with time constants matching those of the generating process improves performance. Alternatively, there could be a general optimal choice for 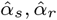 that is independent of the time scale of the generated data, or the time constants could be completely irrelevant and compensated by the complex recurrent dynamics of the network.

The lowest loss is found for values of time constants around the values that were used to generate the data (Fig. 2). This region could differ strongly from the commonly used approximation where either one time constant is ignored (dashed lines) or both time constants are ignored (blue circle). The shape of the loss landscapes changed according to the values of the target time constants, thus indicating that there is not a single choice of time constants that is optimal for any kind of data, irrespective of the underlying generative process.

**Figure 2:**
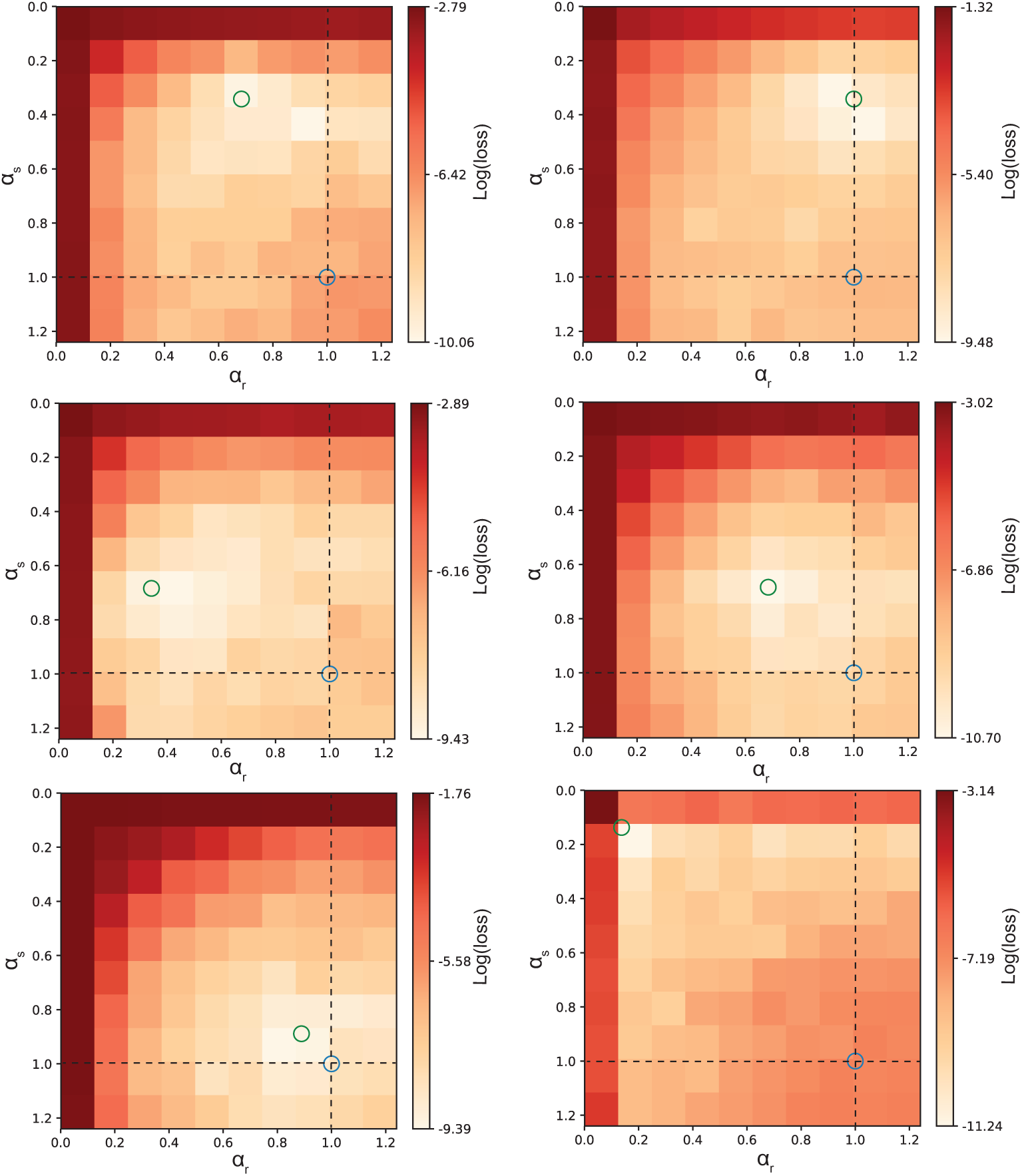
Grid search reveals optimal time constants improve performance. Data were generated for six different combinations of target time constants. A grid search was performed for each of these target pairs, as shown in the different panels (the target time constants are marked by the green circle in every panel). The dashed lines indicate the approximation where either 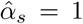 or 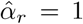. The intersection of both dashed lines indicates the Elman solution (blue circle). The region of lower loss for all generated data examples centers around the actual values (*α*_*s*_, *α*_*r*_) used to generate the data. This indicates that there is a performance benefit from choosing the correct time constants that is not compensated for by the recurrent interactions between neurons.

These results show that the network can recover the time constants of the underlying process that generated certain data. Despite the rich dynamics that RNNs can develop, this is not able to fully compensate for a choice of time constants that is different from the optimal one. At the same time we do not find a symmetric solution (symmetric in the sense that exchanging 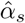 and 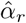 results in the same performance after training), which means that 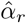 and 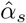 themselves cannot compensate for each other. Both time constants are thus important and an approximation with a single time constant will lead to sub-optimal solutions.

### Learning optimal time constants through backpropagation

From the previous results it becomes clear that choosing the right combination of time constants is beneficial to the performance of an RNN. Most of the time we do not know the underlying time scales of our data though. For this reason it could be useful to let the model learn the time constants via BPTT.^3^

The learning trajectories of the time constants for two example choices of time constants are shown in Fig. 3A, where the learning trajectories of 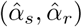 is plotted upon the loss landscape of the same simulations as in Fig. 2 (top and middle left panels, i.e. (*α*_*s*_ = 0.34, *α*_*r*_ = 0.68) and (*α*_*s*_ = 0.68, *α*_*r*_ = 0.34)). Independent of the initialization of the time constants, they clearly converge towards the region of lowest loss obtained from the grid search experiment and recovers the the time constants used in the generative process. Thus the time constants are identifiable and can be learned effectively through BPTT. This means that we can infer information about the time scale of the data by training models to predict dynamic responses from a set of input stimuli. At the same time, the adaptive time constants significantly improved performance over the standard Elman units (Fig. 3B), showing the importance of these time constants.

**Figure 3:**
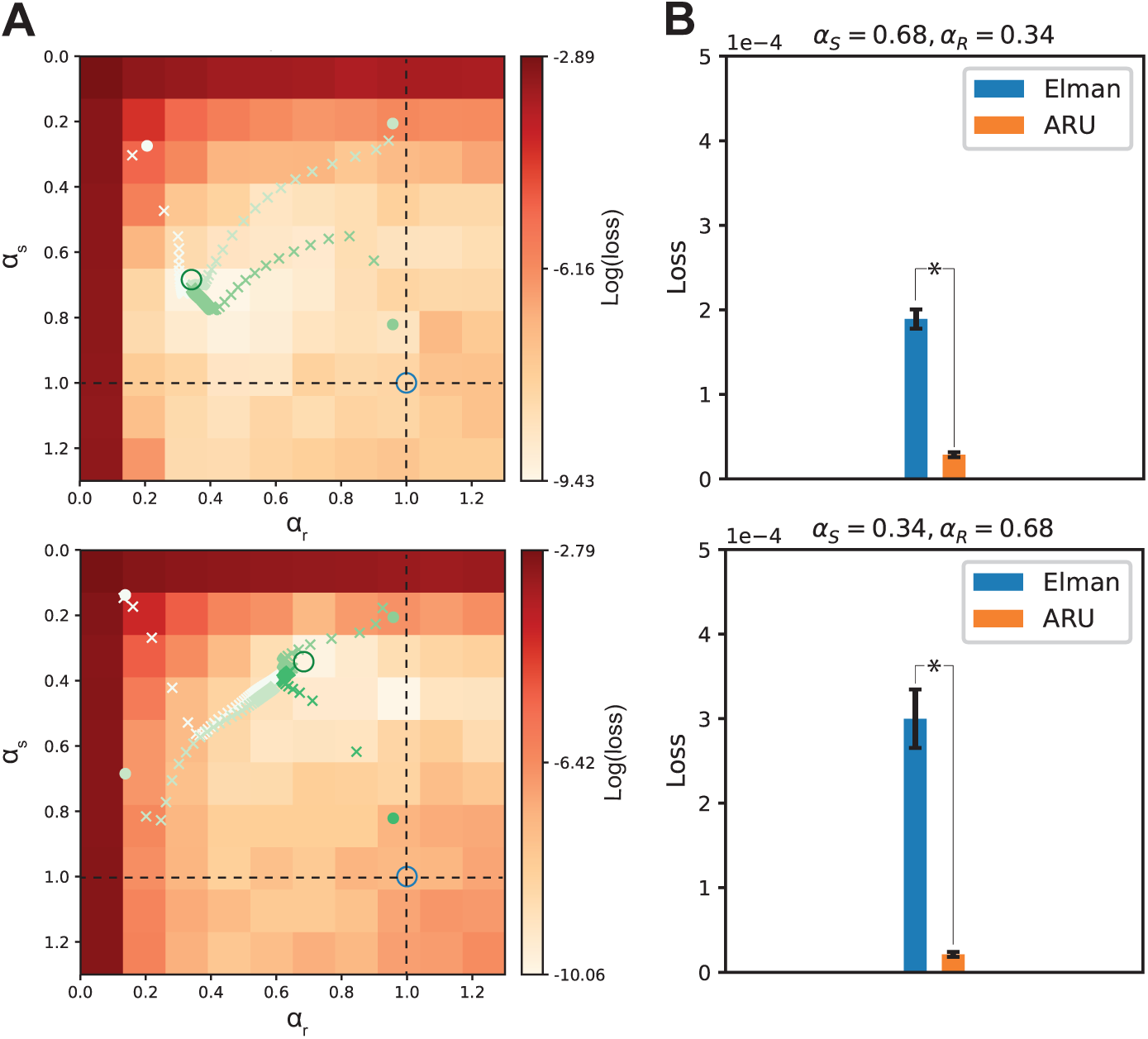
Optimizing time constants through backpropagation. Instead of choosing the time constants manually, the BPTT algorithm was used to optimize them, along with the rest of the parameters of the network. **(A)** For two combinations of target time constants (*α*_*s*_ = 0.684, *α*_*r*_ = 0.34, top panel) and (*α*_*s*_ = 0.34, *α*_*r*_ = 0.68, bottom panel) the optimization of the time constants is plotted over the grid search results of Fig. 2. The target time constants are marked by a green circle. Different initializations of the time constants were tested, as indicated by the filled circles with different shades of green. The learned values after each epoch are marked by a cross of the same color. Independent of initialization, the learned time constants all converge to the target time constants, closely recovering the correct values. **(B)** The performance of the ARU model with learnable time constants was compared with the classical Elman model. The ARU model performed significantly better than the Elman network (\**p <* 1 · 10^−11^ and \**p <* 1 · 10^−6^ for the top and bottom panels, respectively) for both data sets (error bars represent standard error over 20 repetitions).

### Influence of network size on time scale recovery

Simple Elman networks can be seen as universal approximators when given enough hidden units [27]. Therefore, it is possible that a benefit in performance and the recovery of the actual time constants is less in larger networks. We tested networks with different numbers of hidden units in the trained model, as shown in Fig. 4. From these results it is clear that more hidden units do not make the time constants less important with respect to model adaptability: the region of lower loss around the optimal point does not expand as the number of hidden units increases. When learning the time constants, the precision of the recovery does not change with network size either (Fig. 4, blue crosses). Despite the fact that larger networks could learn more complex dynamics, possibly overcoming a non-optimal choice of time constants, the ability to effectively recover time constants of the underlying dynamical process remains. This is promising for future applications on more complex dynamical problems requiring large-scale networks, for example for explaining neural responses in the brain.

**Figure 4:**
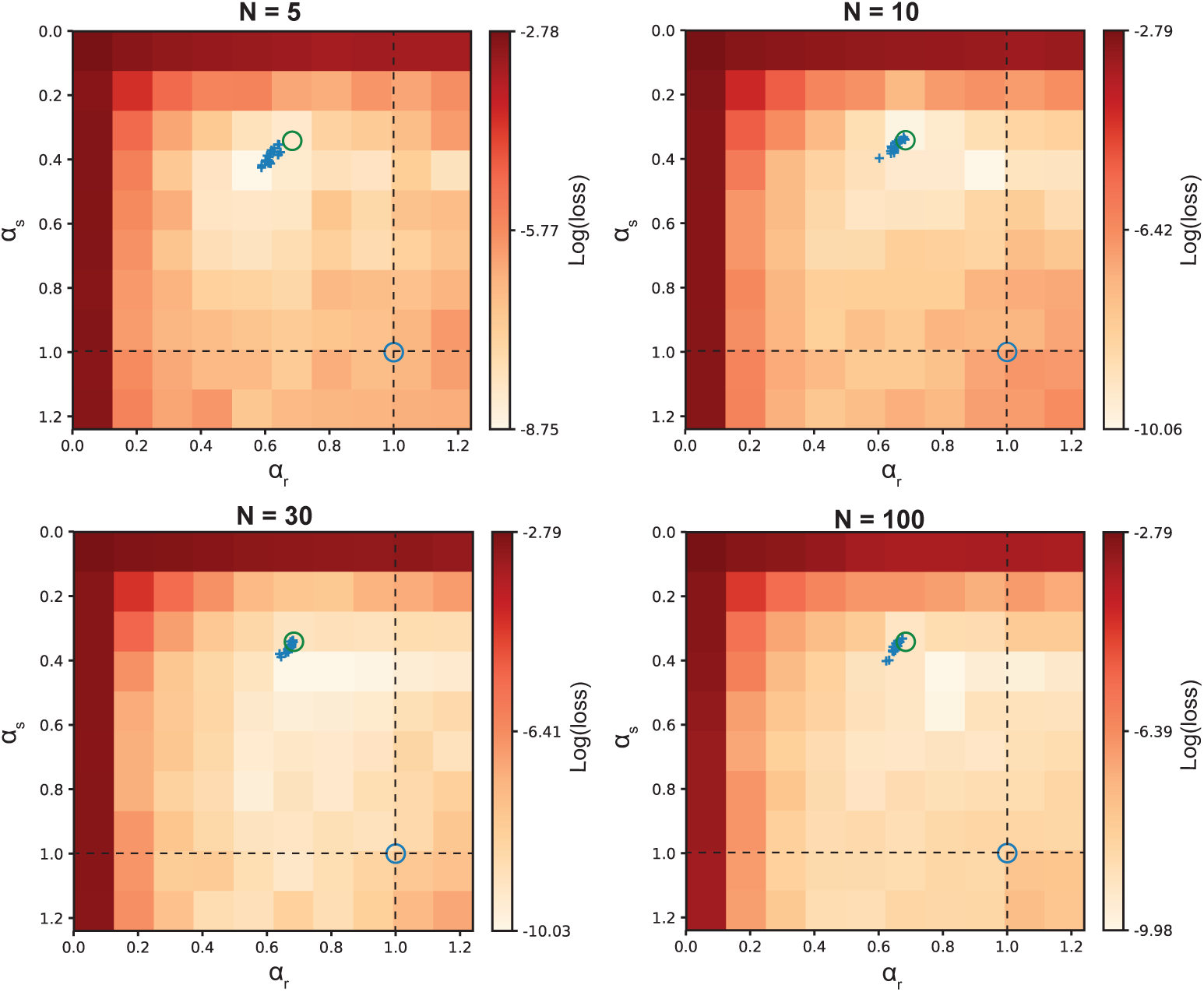
Effect of network size on recovering time parameters. The number of hidden units in the trained model *N* was varied to test the effect on the loss landscape. The target model that generated the data of these simulations was equipped with time constants (*α*_*s*_ = 0.34, *α*_*r*_ = 0.68) as indicated with the green circle, and 10 hidden units. Networks with 5, 10, 30 and 100 hidden units were trained on the generated data. Both a grid search over fixed time constants was performed and networks with adaptive time constants were trained with their final learned time constants indicated as blue crosses (20 repetitions). A larger network size did not decrease the ability of the network to recover the underlying target time constants.

### Time scale recovery depends on activation function

A diverse set of activation functions can be used for the nonlinearities in RNNs. Our previous simulations were done with the commonly used sigmoid activation function. Since it is possible that the recovery of the correct time constants depends on the choice of activation function for the network, we also tested another commonly used activation functions, namely the rectified linear unit (ReLU) (Fig. 5).

**Figure 5:**
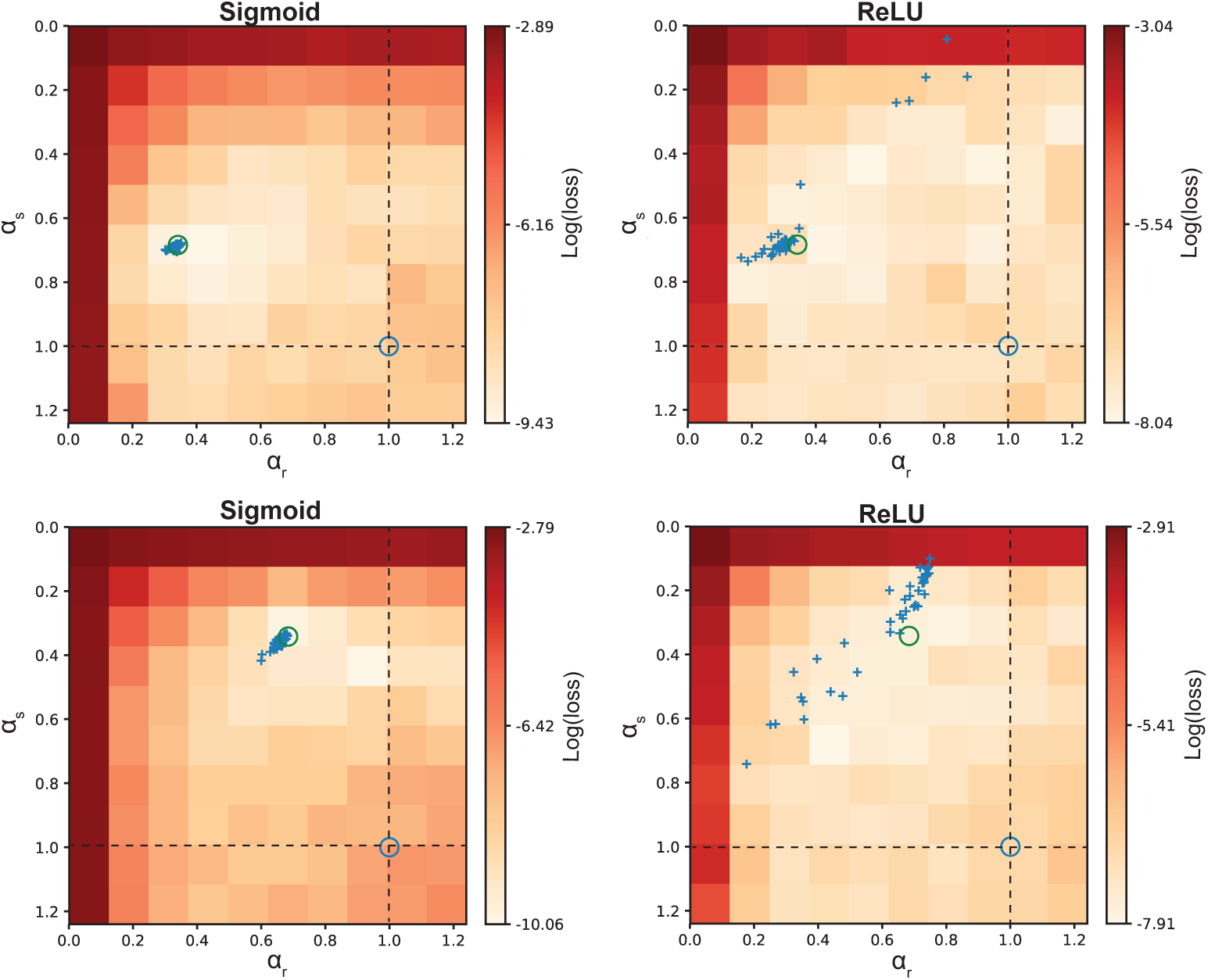
Effect of activation function on recovering time parematers. Networks were either equipped with a sigmoid activation function (left) or a ReLU activation function (right). For two combinations of target time constants (*α*_*s*_ = 0.68, *α*_*r*_ = 0.34, upper panels) and (*α*_*s*_ = 0.34, *α*_*r*_ = 0.68, lower panels) a grid search was performed over fixed time constants and networks with adaptive time constants were trained with their final learned time constants indicated as blue crosses (40 repetitions). Target time constants are indicated with a green circle. The networks with a sigmoid activation function have a region of lowest loss contained around the target values. However networks with an ReLU activation function have a much wider basin of time constants associated with minimal loss. Learned time constants do not recover the target time constants for the ReLU activation function uniquely but form a band symmetric around the diagonal *α*_*s*_ = 0, *α*_*r*_ = 0. The grid search loss region for the ReLU activation function is also symmetric around the diagonal, indicating an interchangeability of time constant 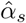 and 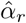. This could be the result of the interchangeability in the linear regime of the ReLU activation function (see Supplementary materials).

While the loss landscape clearly shows a region of lower loss around the time constants used to generate the data in the case of sigmoid activation functions, the loss landscape becomes more shallow and wider in the case of ReLU activation functions, making it harder to identify a unique combination of time constants that leads to the lowest loss. The loss region also becomes more symmetric with respect to the diagonal 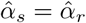. An explanation for this can be that a linear activation function (when in the linear regime) makes the two dynamical processes of each unit (eq. 8, 9) reducible to one, in which we have new time constants which are invariant for exchange of 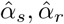 (see Supplementary materials for a derivation). When optimizing the time constants with BPTT, we also find that there is no good recovery of the original time constants (Fig. 5, blue crosses). If our goal is thus to recover the time constants governing the data at hand, the ReLU activation function is not suitable and the sigmoid activation function is a better option.

### Learning a range of time scales

In the previous sections, the learned time constants were shared over all neurons in the network. While this might be sufficient to describe processes with a single underlying pair of time constants, it might not be optimal to describe processes that are governed by a whole range of time scales. To enhance the expressiveness of the network, we learn individual time constants per neuron. Besides increasing performance it could possibly provide insight in the range of time scales underlying the data, where data with a single underlying time scale would lead to narrow range of learnt time constants, and data with multiple different underlying time scales would lead to a broad range of learned time constants.

To test whether such a network that can learn individual time constants per neuron is informative about the range of time scales underlying a process, we generated data with different distributions of time constants drawn from a truncated Gaussian distribution. Subsequently four different neural network models were trained. A network with global adaptive time constants and a network with local adaptive time constants per unit were compared against a standard Elman network and a network composed of the frequently used gated recurrent units (GRU’s) [28].

The learned distribution of time constants was compared with the distribution of time constants used to generate the data. Figure 6A shows the standard deviation of the learned time constants versus the standard deviation of time constants used to generate three different datasets. The network clearly learns to adjust its time constants to the underlying distribution of the data. The range of time constants is recovered such that it can inform us about the underlying time constant distribution. At the same time the network with individual time constants outperforms the network without individual time constants, the Elman network and the GRU network (Fig. 6B). The largest performance difference is found for the dataset generated with the broadest range of time constants, demonstrating the benefit of learning individual time constants on datasets with different underlying processes. While the GRU network has three times as many parameters it performs worse than the network with individual time constants, showing the importance of a network that is architecturally similar to the process it tries to model, making a case for such biologically plausible networks when modelling neural processes.

**Figure 6:**
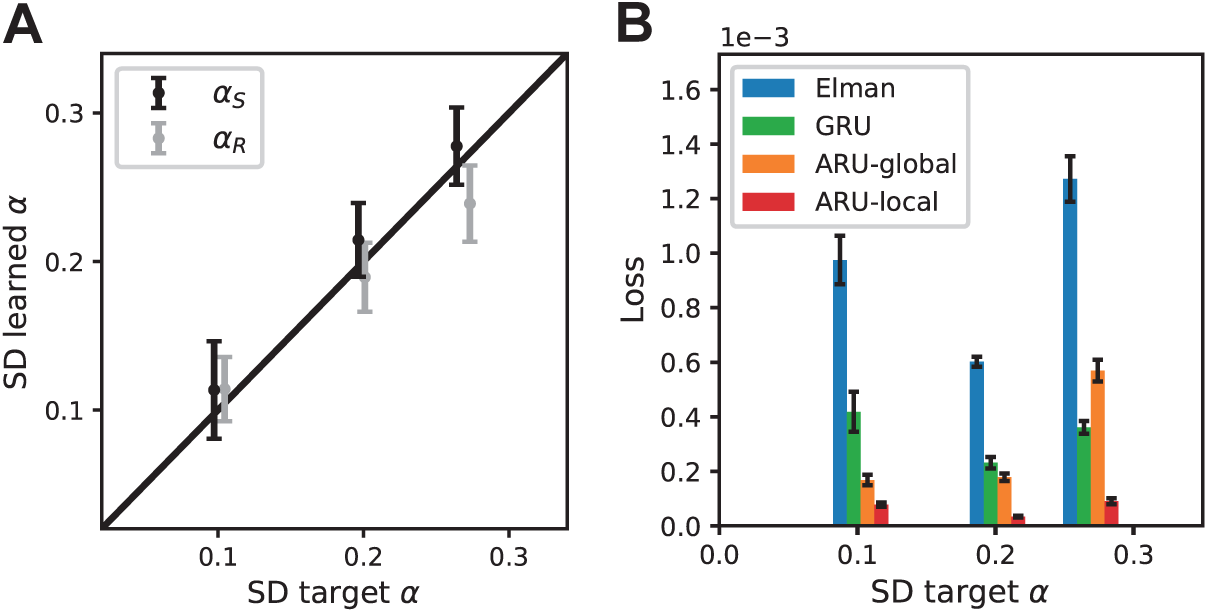
Recovering a distribution of time constants. Data was generated for three time constant distributions with increasing standard deviation (SD). **(A)** Networks trained on this data adopted a range of time constants with similar SD. **(B)** The model with local time constants performs best on the data, improving over the model with global time constants, especially on data generated from a distribution with high SD (error bars represent standard error over 20 repetitions).

### Adaptive time scales increase memory capacity

Adaptive time scales can lead to slower dynamics in a trained network. This should result in an enhanced capacity to retain memory over longer time scales. To test this idea we designed a simple memory task where the memory length could be varied [29]. In this simple task networks had to remember the input that the network received *N* time steps back, with *N* ∈ {5, 10, 20, 30, 40, 50}. The longer back in time the network had to remember its input, the more difficult the task became, demanding enhanced memory capacity. We compared the standard Elman network and GRU network, with a network with a pair of global adaptive time constants, and a network with local adaptive time constants for every unit. Figure 7A shows the performance of the networks on the different memory lengths. The standard Elman network is no longer able to learn the task for retention periods beyond 10 steps, while both networks with adaptive time constants learn better than chance level until memory lengths of 40 steps. The memory capacity of the GRU network is even larger, but this network uses three times as many parameters. To investigate how the adaptive time constants played a role in this increased memory capacity, we averaged the time constants over successfully trained models for the different memory lengths. The learned time constants decreases for longer memory lengths for both network models (Fig. 7B,C), indicating that the slower internal dynamics helped maintain memories over longer time windows.

**Figure 7:**
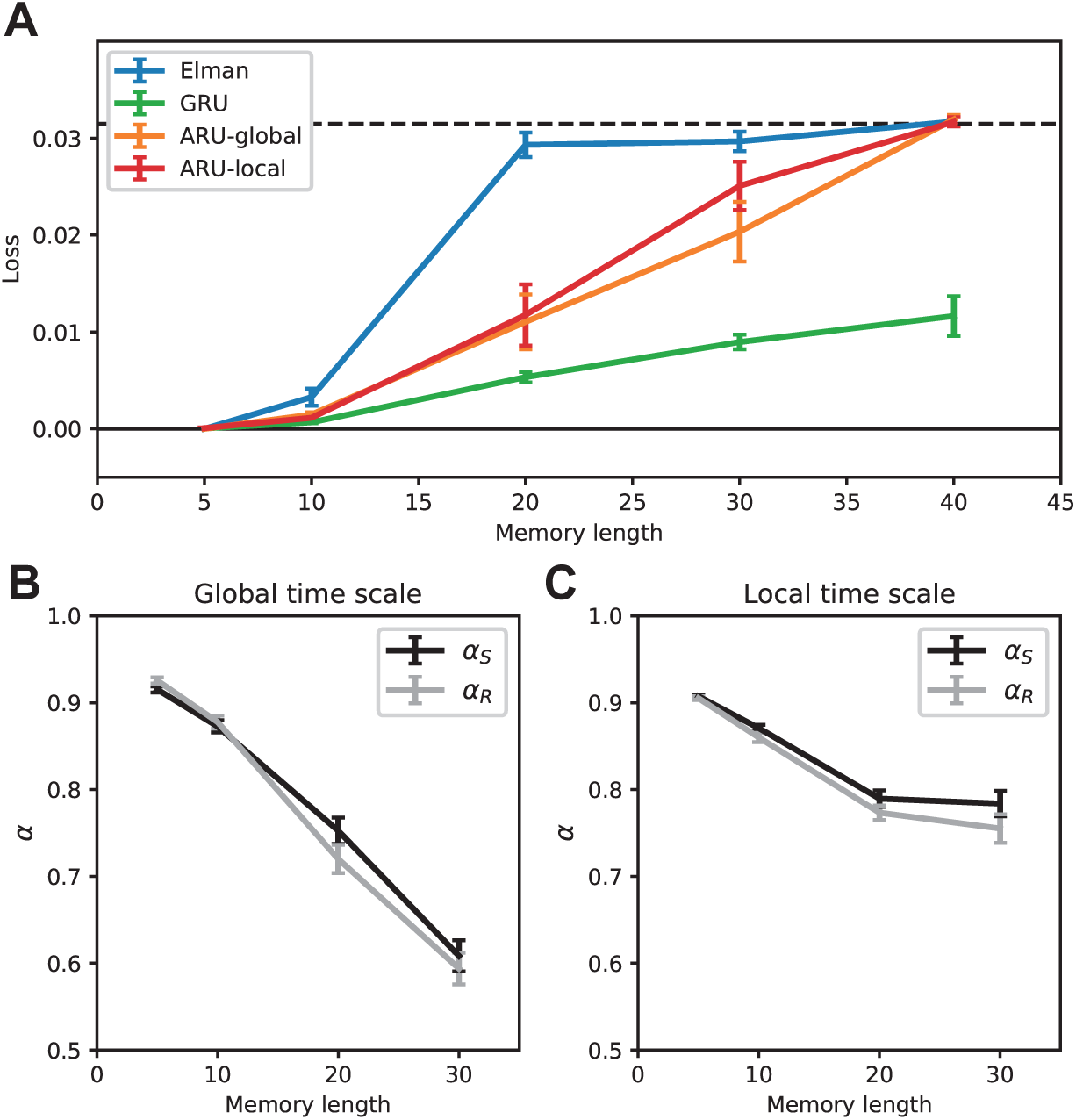
Adaptive time scales increase memory capacity. Three different model variations were tested on a memory capacity task. The number of time steps the input had to be remembered varied from 5 to 40. **(A)** The model using standard Elman units had the lowest memory capacity, performing better than chance up to memory lengths of 10 time steps. Both the model with global and local time constants performed better than chance up to memory lengths of 30 time steps (dashed line represents chance level, which is defined as predicting the average over all time steps). **(B**,**C)** The models with adaptive time constants learned slower time constants with increasing memory lengths, increasing the capacity to maintain memories over longer time scales (error bars represent standard error over 20 repetitions).

### A hierarchy of time scales

Recent findings have shown a hierarchy of time scales in the visual cortex with lower visual areas responding at faster time scales to changes in visual input and higher visual areas responding at slower time scales [7]. This raises the question whether such a temporal hierarchy also emerges when we stack multiple layers of ARU units, each able to adapt its own time constant, on top of each other. To test this, data with a combination of a fast (10 Hz) and slow (2 Hz) sinusoidal signal was generated. A two-layer recurrent network was created consisting of ARU units with feedforward and feedback connections between both layers (Fig. 8A). As a task, the first layer of the network had to predict the next time step of the signal given the current time step of the signal. During learning the two layers of the network specialized their time constants differently to optimize the prediction of the signal (Fig. 8C). After successfully learning the task the network had developed a hierarchy of time constants, with the first layer having a faster time constant responding to quick changes in the signal and the second layer having a slower time constant, responding to slower changes in the signal (Fig. 8B).

**Figure 8:**
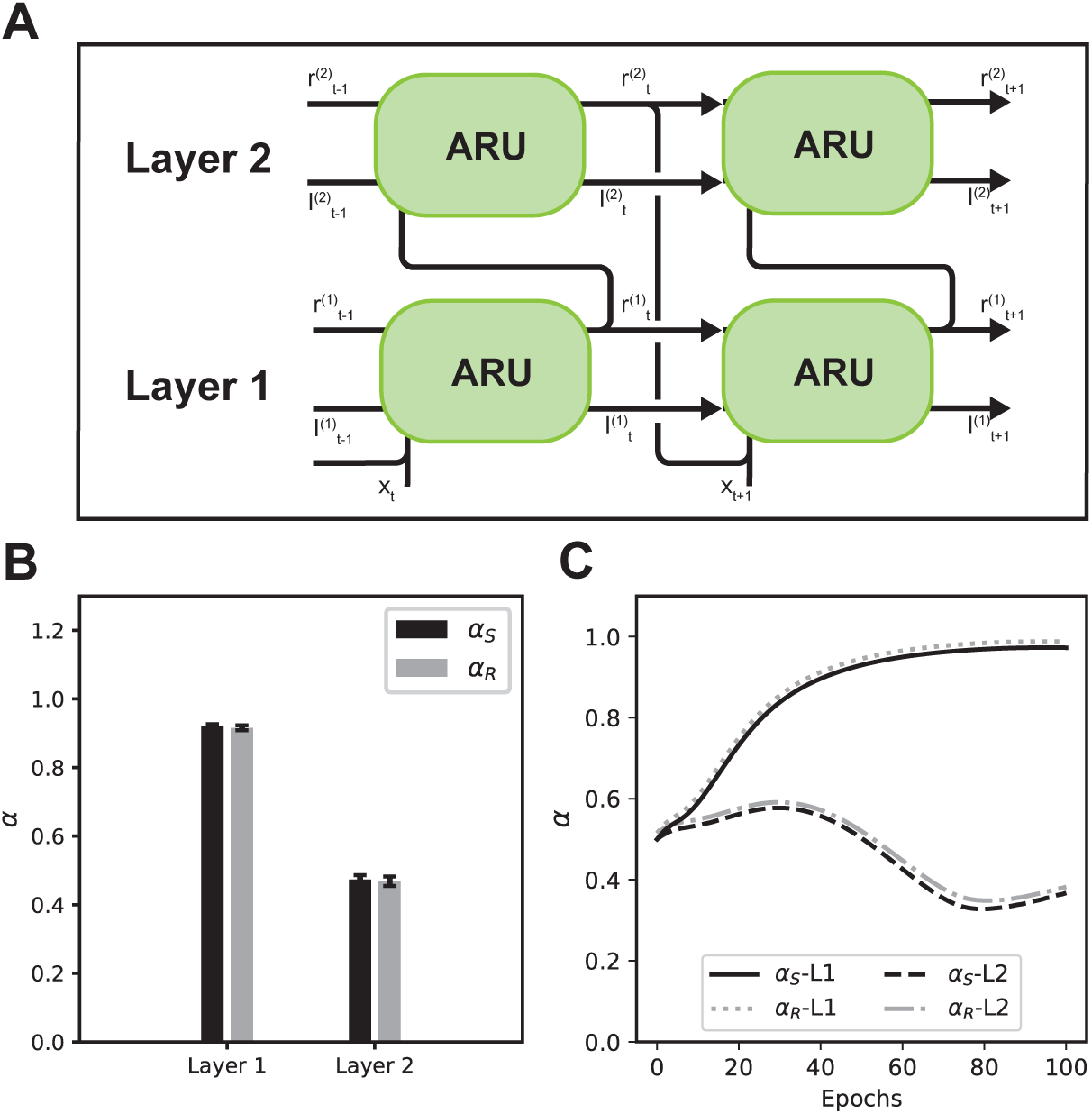
Learning a hierarchy of time scales. **(A)** A two-layer recurrent neural network was created, consisting of ARUs with a global time constant per layer. Both layers were connected through feedfoward and feedback connectivity. **(B)** A temporal hierarchy emerged with the time constants learned by the first supporting much faster dynamics than the time constants learned by the second layer (error bars represent standard error over 20 repetitions). **(C)** Example of learning trajectory of time constants in both layers over 100 training epochs (trajectories for *α*_*R*_ have been offset by 0.01 for visibility).

## Discussion

### Summary of results

We showed that rate-based RNN models can improve performance by including adaptive time constants. In particular, we showed that a particular choice of the time constants can in general increase performance with respect to the commonly used approximations where *α*_*s*_, *α*_*r*_ or both are set to 1. Furthermore, we showed that these time constants can be learned efficiently via the BPTT algorithm. The learned time constants recover the time scales underlying the original process, thus giving valuable insight in the temporal structure of the data. The activation function plays an important role in recovering the time scales. While a sigmoid activation function leads to recovery of both time scales, this is not the case for the ReLU activation function. The recovery of the time scales is independent of the number of hidden units in the network. Although a larger network is theoretically more flexible to compensate for suboptimal choices of time constants, this does not change the ability of the network to recover the correct values. When the dynamics of a data set are governed by multiple time scales, the spread in time constants is successfully recovered by a network where every unit can learn an individual time constant. Since slower time constants can lead to longer memory retention, we tested the ability of the networks to maintain memories over longer periods. The adaptive time constants led to an increased memory capacity compared to the standard Elman approximation. Two layers of ARU units were stacked together to investigate whether a hierarchy of time constants is learned from data composed of multiple time scales. Indeed we find that the first layer of the network learned to respond at a fast time scale, while the second layer learned to respond at a slow time scale, opening avenues to investigate such temporal hierarchies using RNNs.

### Relation to previous work

These results build on previous attempts to enhance RNNs by exploiting the time scale of the dynamics of the data. Adaptive time constants have been proposed in continuous neural networks as a way to increase the expressiveness of an RNN [21]. In this work a purely theoretical derivation was provided for learning time constants and time delays of the continuous neural network units. Other work showed that equipping neurons in a network with different fixed time constants, could be beneficial on tasks having both long-term and short-term temporal relations [30]. In this setting, however, time constants could not be effectively estimated through training. Similarly, different fixed time constants improved performance for recurrent spiking neural networks, approaching the performance of long short-term memory (LSTM) units [31]. More recent work has shown that, by stacking multiple RNN layers with progressively slower fixed time constants, better performance was achieved on predicting motor patterns [32]. It has also been shown that adjusting time constants by numerical integration outperformed the adjustment of time delays in the context of a chaotic process [22]. However, the computational cost of numerical integration hampered the extension to large-scale networks and real world problems. More recent work has attempted to create modules in an RNN, each with its own fixed temporal preference, to improve learning dynamical data with different temporal dependencies [33]. However, these time scales are fixed by the architecture’s connections and can not be optimized during training. Here we implemented the learning of time constants using BPTT, with the benefit of easily extending the implementation of such time scales to more complex network architectures.

Studies into the role of time scales in the brain have revealed a temporal hierarchy, much like the spatial hierarchies found in the visual cortex [7]. Similar to the spatial hierarchy exhibited by the receptive fields of neurons in the visual cortex, there is also a temporal hierarchy of neurons responding to fast stimulus changes in early visual cortex, while neurons in higher visual cortex respond to slower changes. A tempting explanation for the emergence of such a hierarchy is that the hierarchical causal structure of the outside world shapes the representations of the brain [10]. Neural networks have been used to study the emergence of a temporal hierarchy in producing motor patterns [32], showing a functional role of higher layers with slow dynamics composing different motor primitives together in the lower layers with fast dynamics. However, such hierarchies were partially imposed by fixing the time scales of subsequent layers to progressively slower time scales. Here we show that when time scales are learned, such a temporal hierarchy emerges automatically from the data, without the need for imposing any architectural constraints. These results provide promising avenues for investigating the emergence and functioning of temporal hierarchies in the brain.

### Benefits of learning time constants

Learning time constants can be useful when modeling neuroscientific data. To gain deeper insight in the functional role of the temporal hierarchy in the visual cortex, neural networks with emerging temporal hierarchies may be used as encoding models, similar to how spatial hierarchies were successfully modelled by feedforward convolutional networks [34]. By training RNNs with adaptive time scales to predict neural responses we can combine knowledge about the temporal structure of the cortex with insight in the actual information that is processed by a neural population. Questions regarding what properties of the data lead to the emergence of a temporal hierarchy, and how task requirements can influence the time scales which certain cortical areas are responsive to can be investigated through such models and validated against experimental data.

Another example stems from the motor control literature. The motor patterns formed by the brain are thought to be composed of smaller motor ‘primitives’ [18]. These primitives can be combined in a temporal sequence by a hierarchical process, where a slower process composes these primitives in meaningful motor patterns. Again, our approach could gain insight in which parts of the motor cortex represent these slower processes and which parts represent the faster primitives processes. Our approach can help us gain valuable insight in the functional relevance of the many dynamical processes in the brain.

Besides modeling neuroscientific data, learning time constants can also be useful in machine learning applications. Building intelligent models to learn complex dynamical tasks, could benefit from artificial neurons that can learn and adapt to the time scales relevant for the problem at hand. Similar to the brain, artificial systems are often trained on natural data, where processes evolve over different time scales and certain information remains relevant over longer time scales while other information is only relevant for a very short period of time. An interesting application is the generation of natural movements for robots. It has been shown that using different time scales in a hierarchical model improves the learned motor patterns [32, 35, 18]. Similarly, the recognition of actions performed by humans from video data has been shown to benefit from such an temporal hierarchical structure [36, 37]. However, setting such a hierarchy of time scales by hand is cumbersome and does not guarantee optimal results. Including the learning of these time scales in the optimization of the network ensures automatic optimization and could lead to the automatic emergence of models with relevant hierarchical time scales.

### Future work

Our approach enables researchers to build more expressive neural network models, and at the same time recover relevant temporal information from the data. Building towards large-scale models to predict brain responses over a large number of areas will gain us valuable insight in the dynamics of brain processes. Extensions towards learning time delays between brain areas or learning time constants that can be modulated by the input are promising future steps. Keeping such complex models well-behaved during optimization, without falling prey to local optima, is an important challenge for future work. On the other hand there is the challenge of keeping networks interpretable by letting the network learn relevant biological parameters (such as time constants or time delays). Further work is needed to investigate how well such parameters can be recovered in large and complex networks where such complex dynamics might emerge that learning the correct parameters can be circumvented [38].

### Conclusions

We found that making standard RNN models more biologically plausible by introducing learnable time constants improves performance of the model and increases its memory capacity, while at the same time enabling us to recover the time scales of the underlying processes from the data. Gaining explicit knowledge about the time scales at which processes unfold can improve our understanding of hierarchical temporal dynamics in the brain. At the same time, facilitate explicit time scales the creation of more expressive and interpretable machine learning models, which shows that embracing principles of neural computation can help us to develop more powerful AI systems [39].

## Acknowledgements

This research was supported by VIDI grant number 639.072.513 of The Netherlands Organization for Scientific Research (NWO).

## Supplementary materials

### Derivative of synaptic current

The derivative of the synaptic current can be calculated using the Leibniz integral rule. Starting from Eq. 6 and using the exponential kernel 3 we obtain

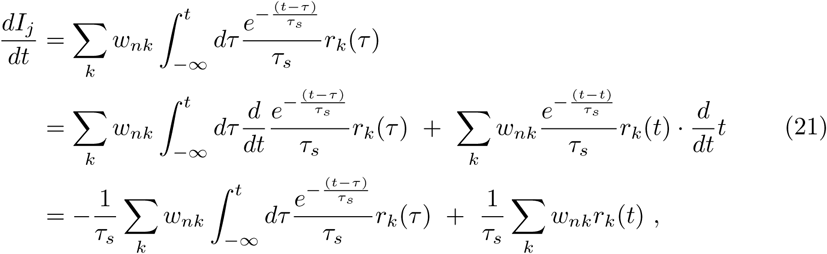

### Linear activation function and symmetric time constants

To understand why ReLU activation functions do not recover the time parameters well it could be informative to consider what happens when we make a linear approximation of the activation function. We start again with Eqs. (8) and (9):

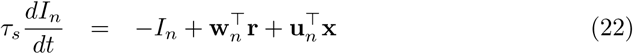

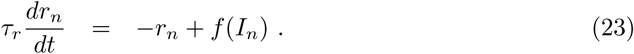

We use a linear activation function, that is *f* (*I*) = *I*. Rewriting Eq. 23

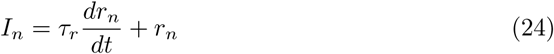

and computing the time derivative

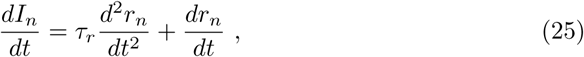

it is possible to substitute expressions for *I*_*n*_ and 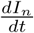 into Eq. 22 to get

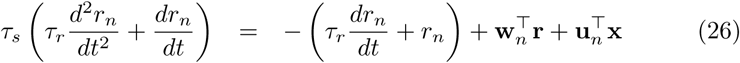

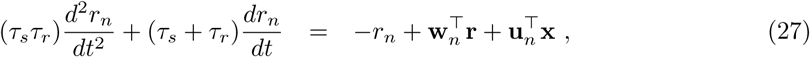

which is a single second-order differential equation, describing the firing rate dynamics only in terms of firing rates. We have two new time constants that govern the dynamics, *τ*_1_ = (*τ*_*s*_ + *τ*_*r*_) and *τ*_2_ = (*τ*_*s*_*τ*_*r*_), which are invariant under exchange of *τ*_*s*_, *τ*_*r*_, and therefore symmetric. This explains why the time constants are not uniquely recoverable for ReLU units, but instead are exchangeable for one another.

## Footnotes

1 In what follows, we will refer to *α*_*r*_, *α*_*s*_ as ‘time constants’ as well, keeping in mind their relation to *τ*_*s*_ and *τ*_*r*_. Time constants expressed as *α* are just more practical for our purpose.

2 Values for 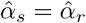 higher than one indicate a negative correlation between subsequent time steps, which is not biologically plausible but were included out of exploratory interest.

3 We investigated whether these parameters can be inferred with gradient descent algorithms. Similar results hold for Stochastic Gradient Descent and Adam optimizer [26]).

